# Detergent induced structural perturbations in peanut agglutinin: Insights from spectroscopic and molecular dynamic simulation studies

**DOI:** 10.1101/2023.07.03.547500

**Authors:** Shreyasi Asthana, Sonali Mohanty, Harshit Kalra, Nandini Karunanethi, Sujit Kumar, Nikhil Agrawal, Suman Jha

**Affiliations:** Department of Life Science, National Institute of Technology, Rourkela, Odisha-769008, India; Latvian Institute of Organic Synthesis, Aizkraukles 21, LV, Riga-1006, Latvia; College of Health Sciences, University of KwaZulu-Natal, Durban-4000, South Africa; Centre for Proteomics and Drug Discovery, Amity Univerity Maharastra, Mumbai-410206, Maharastra, India

**Keywords:** Peanut agglutinin, Sodium dodecyl sulphate, Cetyltrimethylammonium bromide, Molecular docking, Molecular dynamic simulation

## Abstract

The three dimensional structure of a protein is very important for its structure. Studies relating to protein structure have been numerous and the effect of denaturants on proteins can help understand the process of protein folding and misfolding. Detergents are important denaturants and play important roles in various fields. Here we explored the effect of sodium dodecyl sulphate (SDS) and cetyltrimethylammonium bromide (CTAB) on the structure of peanut agglutinin (PNA). The protein was purified from its natural source and impact of SDS and CTAB was studied by circular dichroism, intrinsic fluorescence, 8-anilino-1-napthalenesulfonic acid, molecular docking and molecular dynamics simulation. Pure peanut agglutinin showed a trough at 220 nm and positive ellipticity peak at 195 nm, specific for lectins. Results from the experimental and simulation studies suggest how oppositely charged detergents can interact differently and lead to varied structural perturbations in PNA. Both the surfactants induce all α protein-like circular dichroism in the protein, above its critical micelle concentrations, with significant change in accessible surface area that became more hydrophobic upon the treatment. Major interactions between the surfactants and protein, resulting in PNA conformational rearrangement, are electrostatic and van der Waals interactions. However, CTAB, like a cationic surfactant, has similar effects as anionic surfactant (SDS) but at significantly very low concentration. Though the effects followed same pattern in both the surfactant treatment, i.e. above respective CMC, the surfactants were inducing α-helix/coil like conformation in PNA.

## 1. INTRODUCTION

The three-dimensional structure of a protein is very essential in determining its stability, specificity, and activity. Therefore, understanding the pathways a protein follows during its folding and unfolding is essential to solve the protein folding problem[1-3]. Studies on protein folding and misfolding provide an idea about the pathways and mechanisms that a protein may take up during its folding process[4]. Perturbations in a protein structure can be brought about by many physical and chemical factors[5-7]. One of the most common chemical denaturants used for protein unfolding studies is Guanidium Hydrochloride (GdnHCl). The precise mechanism of its effect on protein structure is yet to be deciphered[8]. The other most important physical factor used to study protein denaturation is temperature. At high temperatures, proteins tend to lose their three-dimensional structure[9]. However, the mechanism of unfolding is different in both the cases. Apart from these denaturants, other most common agents routinely used to disrupt protein structures, in several experimental procedures, are detergents.

Nonetheless, studies relating to the denaturation of tertiary/quaternary proteins by detergents have been rare and consequently, their mechanism of action is also not very clear[10-12]. Sodium Dodecyl Sulfate (SDS) is an anionic detergent, with a 12-carbon hydrophobic chain. There have been early studies on the binding of SDS with proteins, and the nature of the SDS-protein complex has been elucidated. It was shown that 1 gram of protein can bind to as much as 1.5-2 grams of SDS[13]. Early investigations suggested that unfolded proteins wrap themselves around detergent micelles[14]. Whereas, recent studies suggest tertiary structure unfolding at lower SDS concentrations and chain expansion at high SDS concentration[15]. On the contrary, Cetyl Trimethyl Ammonium Bromide (CTAB), a cationic detergent with a 16-carbon long hydrophobic chain, binding to protein is generally endothermic in nature[16]. However, unfolding is brought about at a higher concentration of CTAB. The shape of binding isotherms is similar for both the cationic and anionic detergent[17, 18]. Molecular Dynamics (MD) Simulations studies on the effect of these detergents on protein structures have also been an area of interest lately. A recent MD simulation described the structural changes in transmembrane helices of bacterial rhodopsin upon interaction with SDS [19]. Similarly, another study showed the association and encapsulation of myoglobin in CTAB micelles that led the complex into aggregation [20].

Peanut Agglutinin (PNA) is a homo-tetrameric, non-glycosylated protein, belonging to the legume lectin family. It is also known as anti-T-agglutinin because of its specificity for the T-antigen. Each subunit of PNA is 273 amino acids long with a molecular weight of ∼27 KDa [21]. PNA structure shows that it has a lectin-specific jellyroll tertiary structural folds. A typical jelly roll motif contains a six-stranded antiparallel flat back sheet and a seven-stranded curved front sheet which are held together by a small five-stranded top sheet. It has two hydrophobic cores, one is present between the two large sheets and the other one is between the curved sheet and the loops protruding from the edges of the sheet [22]. Association of the monomeric subunits into the protein quaternary structure occurs in the most unexpected way, giving rise to an open quaternary structure[23]. Subunits 1-2 associate in canonical mode, while 1-4 and 2-3 associate in a back-to-back manner (Fig.1B). Canonical dimerization is mediated through six water bridges, unlike other canonical interfaces. Subunits 3-4 do not interact directly and form a unique “open” quaternary association, thus lacking a 4-fold symmetry [23]. Therefore, the integrity of PNA tertiary structure is prone to subtle variance in local physicochemical conditions to cause divergent modes of oligomerization [24]. Residues in the hydrophobic core are involved in hydrogen bonds or hydrophobic interactions with native ligands, like saccharide and metal ion coordination [25]. GdnHCl-induced unfolding studies have shown that PNA follows a three-state unfolding mechanism, where the molten globule-like intermediate retains its carbohydrate binding coordinates [26]. The present study aims to decipher the conformational dynamics in PNA in presence of anionic and cationic detergents, like SDS and CTAB respectively. To this end, standard spectroscopic techniques have been explored to study the conformational dynamics following different probes, like circular dichroism and steady state fluorescence spectroscopies, whereas docking and MD simulations have been used to underline the interaction interfaces between PNA and the surfactants at the atomic and structural dynamics levels.

## 2. METHODOLOGY

### 2.1 MATERIALS

Sodium Dodecyl Sulphate and 8-anilino-1-napthalenesulfonic acid (ANS) were obtained from Sigma (USA). CTAB, sodium alginate and guar gum were obtained from Himedia. Peanuts were purchased from a local store in Rourkela, and the PNA was purified using a previously described protocol [27]. The PNA purification was established using SDS-PAGE, where a single band of M_wt_ 27.5 KDa was observed. The concentration of purified protein was calculated by taking specific absorbance at 280 nm (A_1%, 1cm_) of 7.7 [21]. All the other reagents used were of analytical grade. All the buffers and solutions were prepared in MilliQ water, and they were filtered through a 0.45 µm syringe filter before measurements. The molar concentration of ANS was determined spectrophotometrically by an extinction coefficient of 7800 M^−1^cm^−1^ at 372 nm [28]. All the experiments were performed in 10 mM phosphate buffer (pH 7.4) at 25 °C.

### 2.2 PROTEIN DENATURATION AND SPECTROSCOPIC MEASUREMENTS

Stock solutions of SDS and CTAB were prepared in 10 mM phosphate buffer (pH 7.4) and filtered through a 0.45 µm membrane filter. For each denaturation experiment, a known amount of phosphate buffer was diluted with a fixed volume of the protein stock solution of desired final concentration, and a varying amount of the concentrated denaturants were mixed to make a final reaction volume of 1 mL. The reaction mixture was further incubated for 8 hrs to attain equilibrium.

Circular Dichroism (CD) spectroscopic measurements were performed in the far UV range (190-260 nm) for 10 µM PNA in the absence and presence of SDS or CTAB in a 1 mm cuvette using JASCO-J1500 CD spectropolarimeter (Tokyo, Japan) purged with N_2_ gas, and equipped with a Jasco Peltier-type temperature controller system maintained at 25 °C. For protein-tryptophan/tyrosine intrinsic fluorescence, 2 µM PNA was incubated with or without the surfactants. Upon the incubation, the fluorescence measurements were done in a 10 mm cuvette, using LS-55 Perkin Elmer Spectrofluorimeter (Waltham, USA). The emission spectra were recorded between 300-400 nm upon excitation at 280 nm with a filter width of 5 nm each, for excitation and emission filters, and at a scan speed of 100 nm/min with increasing concentrations of SDS or CTAB. Similarly, for ANS-based extrinsic fluorescence, 100 µM ANS emission spectra were collected, upon incubation with intact 2 µM PNA or with varying concentrations of the either surfactant-mixed PNA, in the range of 410 – 600 nm after excitation at 388 nm, using filter widths of 5 nm each with a scan rate of 100 nm/min.

### 2.3 MOLECULAR DOCKING AND DYNAMICS SIMULATIONS

AutoDock 4.2 was used to dock single molecules of CTAB and SDS with PNA to predict the interaction pattern between the protein and ligands, as previously mentioned [29]. Briefly, the three-dimensional structures of CTAB and SDS were retrieved from the PubChem database, which is accessible at https://pubchem.ncbi.nlm.nih.gov (CID 5974, 3423265). Using PyMol (Schrödinger), the ligands were converted from 2D structure data file (SDF) to a 3D protein data bank file (PDB) format. Using the AutoDock Tools, bond torsions were assigned to the ligands to prepare them for docking. The 3D coordinates of PNA were also obtained from the RCSB database with PDB ID: 2DV9 (https://www.rcsb.org/structure/2DV9) [30]. Water molecules and heteroatoms were eliminated from the protein structure, whereas hydrogen atoms and gasteiger charges were added to the structure. PNA is a homotetramer, hence docking was done using only chain A. A grid map with the dimensions 126 × 126 × 126 Å^3^ and a spacing of 0.508 Å was created using AutoGrid, and it was centred at 103.855 Å x 45.003 Å x 24.355 Å. Utilizing the Lamarckian Genetic Algorithm (LGA), single ligand molecular docking was carried out. The default values for population size, maximum evaluations, maximum generations, mutation rate, and crossover rate were maintained. Translation, quaternion, and torsion in the docking parameters were likewise left at default. The binding energy represents the sum of minimum energies obtained from 10 docking runs. Final representing figures of the docked complexes were created using PyMol (Schrödinger) and Discovery Studio 2.5 (BIOVIA) for further analysis [31, 32].

To further evaluate the protein conformational dynamics in the presence of surfactants at the residue level, molecular dynamics simulations were run for PNA monomer with respective numbers of the ligands representing critical micelle concentration (CMC) in the defined simulation box. Hence for the molecular dynamics study, PNA monomer (PDB id:2DV9) [33], SDS (PubChem CID: 23700838) and CTAB (PubChem CID:5974)[34] were used. Packmol [35] was used to add molecules of SDS and CTAB in simulation systems, and all the molecules have a minimum cut-off distance, from PNA, 1.0 nm. PNA apo system contained one PNA monomer and 22666 water molecules with 6 Na^+^ ions to neutralize the systems. PNA with an SDS system contained one PNA monomer, 35 SDS and 19388 water molecules with 41 Na^+^ ions to neutralize the systems. PNA with a CTAB system contained one PNA monomer, 7 CTAB, and 19427 water molecules with 7 Br^-^ and 6 Na^+^ ions to neutralize the systems. The number of ligands was calculated employing N = (M/55.5)*W formula, where M is the intended molecular concentration in molar of either the ligand, and W is the number of water molecules in the box. The CHARMM36 force field [36] was used for PNA and the CHARMM general force field [37] was employed for SDS and CTAB, and the TIP3P was used for water [38]. All three systems were energy minimized using the steepest descent algorithm [39], followed by two sequential equilibration simulations using the canonical (NVT) and isobaric−isothermic (NPT) ensemble, and the production simulations were performed using the NPT ensemble for 500 ns for each system. The LINCS algorithm [40] was employed for constraining the bond lengths from heavy atoms to hydrogen, and the SETTLE algorithm [41] was used for water molecule bond length constraints. For long-range electrostatic interactions, the Particle Mesh Ewald (PME) algorithm [42] was used, and for short-range van der Waals (vdW) and columbic interactions, a cut-off distance of 1.2 nm was used. The Parrinello−Rahman algorithm [43] was used for pressure coupling, and the Nose-Hoover algorithm [44] was employed for temperature coupling. All production simulations were performed at a pressure of 1 bar and temperature of 300 K using the GROMACS (version 2016.1) simulation package [45]. GROMACS gmx rms, gmx rmsf, gmx gyrate modules were used for rmsd, rmsf and rog calculations, respectivtely. For RMSD, RMSF and ROG backbone atoms were used, and RMSF was calculated for each residue. DSSP tool [46] was used for the calculation of secondary structure, and gmx mindist tool was used for the calculation of minimum distance between SDS and CTAB molecules from PNA.

## 3. RESULTS AND DISCUSSION

### 3.1 Characteristics of the Native PNA

Peanut Agglutinin (PNA) is a highly stable homotetrameric protein, which showed a lectin-like peculiar circular dichroism (CD) spectra with broad negative ellipticity peak centred at 220 nm and positive ellipticity peak at 195 nm (Fig. 2A). The monomeric subunit of PNA being a jelly roll motif, lacks the fourfold rotational symmetry (Fig 1) [47]. The PNA subunits association, in one interface, is through back β-sheets aligned at an angle of 86° (subunit 1 with 2 and subunit 3 with 4), whereas laterally aligned subunits form two different interfaces called as canonical (between subunit 1 and 4) and open (between subunit 3 and 2) interfaces which makes this lectin-like quaternary structure relatively rigid against surfactant [23, 48]. Although denaturant-mediated unfolding of the quaternary structure is highly unlikely, than the significant conformational rearrangement in the monomeric units [49]. Additionally, PNA is known to exhibit a molten globule-like stable intermediate, when presented to denaturant-based unfolding [48, 49].

**Figure 1:**
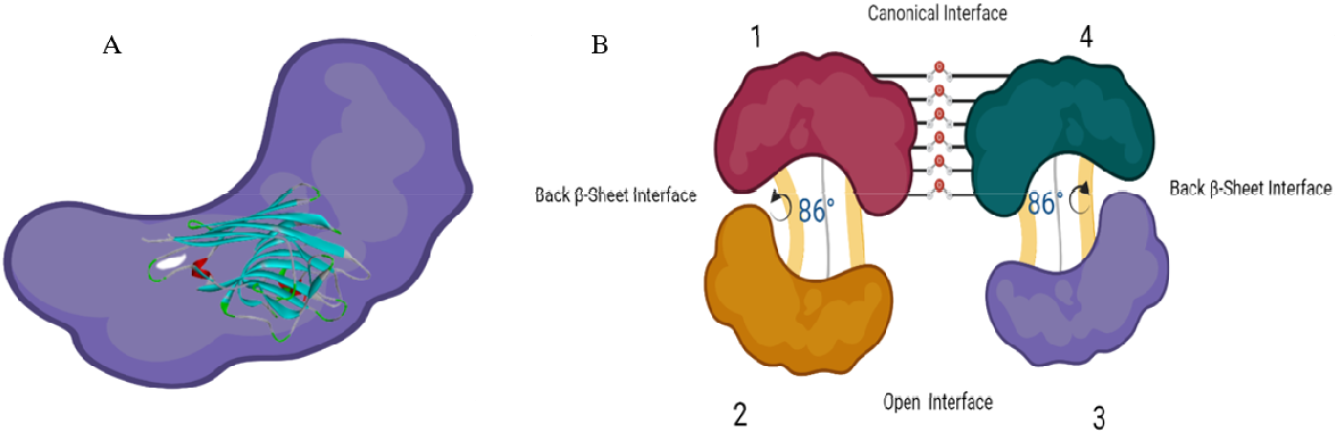
Representative bean shape is equivalent to single monomer of PNA with PDB ID 2DV9 (A). Representative cartoon image of complete PNA (B). The different coloured bean shapes are representation of four PNA monomers. The PNA subunits are associated, in one interface, through back β-sheets aligned at an angle of 86° (between subunit 1 and 2, subunit 3 and 4). Also, laterally aligned subunits form two different interfaces called as canonical (between subunit 1 and 4) and open (between subunit 3 and 2).

**Figure 2:**
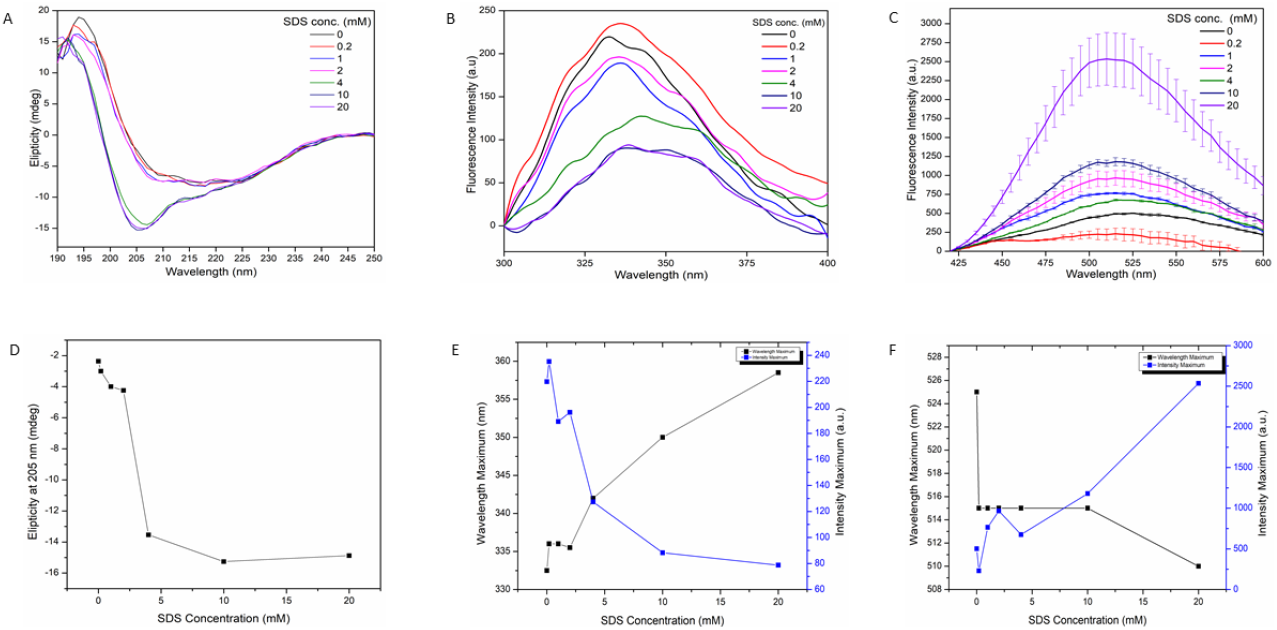
Conformational changes in PNA in presence of varying concentration of SDS as observed using (A) far–UV CD spectra from 190 nm to 250nm; (B) tryptophan fluorescence emission spectra from 300 nm to 400 nm; (C) ANS fluorescence spectra from 420 nm to 600 nm upon binding of PNA at increasing concentration of SDS; (D) ellipticity maxima at 205 nm against increasing concentration of SDS; (E) intensity and wavelength maxima of tryptophan fluorescence; and (F) ANS fluorescence plotted against increasing concentration of SDS.

### 3.2 SDS induced structural perturbations in PNA

Anionic SDS presence, below its CMC, showed an insignificant ellipticity change in PNA with value of -4 and -7 millidegrees (mdeg) at 205 and 220 nm, respectively (Fig. 2A). Interestingly as the concentration of the anionic surfactant was increased above the CMC, PNA underwent a significant conformational change, exhibiting ellipticity corresponding to the all α−protein with traces of β-sheets [48, 50, 51] with negative ellipticity peak and shoulder with value of -15 and -10 mdeg at 205 and 220 nm respectively (Fig. 2A). The abrupt ellipticity change was observed for increasing SDS concentration from 2 mM to 4 mM (Fig. 2D) which interestingly, also represents the phase boundary between monomeric and micellar structure of SDS [52]; CMC of SDS, when dissolved in 10 mM phosphate buffer, is 4.61 ± 0.01 mM [53]. Thus, the conformational changes in PNA is observed at higher SDS concentrations only. In general, considering the change in conformational entropy, random coils/loops are more prone to immediately undergo conformational changes in presence of surfactant than β-sheets or α-helix of a protein [52]. The anticipated interaction interface between SDS and PNA was further validated using single ligand docking of the either surfactant to the protein monomeric subunit (PDB id: 2DV9, Fig. S1), that showed the surfactant binds to loops or relatively dynamic region of PNA (Fig. S2-3). The SDS binding site is present in the second hydrophobic core of the protein subunit, with binding energy of - 3.67 KCal/mol (Table 1). The residues involved in interactions were Trp55, His116, Val142, Lys197, and Pro199 (Fig. S3). The charged sulphate group in SDS, as anticipted, interacted through polar interactions with Lys197 side chain and Val142, whereas Trp55, His116, Lys197 side chains (Table 1) interacted through van der Waals and non-polar interactions with alkyl tail of SDS (Fig. S3). Interestingly, the docking result indicated that, in addition to other interactions, Val142 additionally forms conventional hydrogen bond with the charged SDS head.

**Table 1:** Representing binding energy, interacting residues and bond length of PNA on interaction of SDS and CTAB respectively

| Protein | Ligand | Binding energy | Residues involved | Bond Length |
| --- | --- | --- | --- | --- |
| PNA | SDS | -3.67 Kcal/mol | TRP55, VAL142, HIS116, LYS197, PRO199 | A:VAL142- UNKO:H (2.22302)<br>A:LYS197-UNKO (5.29141)<br>A:PRO199-UNKO (4.44379)<br>A:PRO199- UNKO:C (5.10839)<br>A:TRP55-UNK (5.05277)<br>A:HIS116-UNKO (4.57601) |
|  | CTAB | -2.52 Kcal/mol | THR124,PRO134, ASP136, PRO152 | A:ASP136- UNKO: N (3.2953)<br>A:THR124- UNKO:C (3.13671)<br>A:PRO134- UNKO:C (2.98914)<br>A:ASP136- UNKO:C (2.86272) |
|  |  |  |  | A:PRO152-UNKO (5.29847) |

The involvement of Trp55, was further verified by following the intrinsic fluorescence of PNA-tryptophan and tyrsoine in presence of increasing concentration of SDS (Fig. 2B). Tryptophan or/and tyrosine residues’ fluorescence property change with the change in local physicochemical environment, as the indole ring and 4-hydroxyl phenyl group of tryptophan and tyrosine respectively are hydrophobic in nature and exhibit the fluorescence property that is highly sensitive to the polarity of the accessible medium [54]. In the same line, the intact PNA exhibited fluorescence intensity of 220 a.u. with maxima at 332 nm, upon excitation at 280 nm in the studied conditions (Fig. 2E). Upon addition of SDS, intrinsic fluorescence of PNA decreased from 230 a.u. to 80 a.u. with its increasing concentration (from 0.05 mM to 10 mM), and the emission maxima shifted from 336 nm to 360 nm. Hence, the fluorescence intensity exponentially decreased with significant red shift in emission maxima upon addition of varying SDS concentration to the protein solution. This decrease in the intensity of fluorescence emission, in presence of SDS, indicates that either the local physicochemical conditions around tryptophan/tyrosine residue(s) change towards net higher polarity [48] or progressive presence (aggregation) of hydrophobic functional groups are quenching the fluorescence [55], or both. Hence, the intrinsic fluorescence indicates that, if not all tryptophan and tyrosine of PNA, atleast Trp55, as it is directly involved in interaction with the SDS, and Trp153, for being in proximity to SDS binding pocket, are getting further either burried into hydrophobic environment resulting in fluorescence quenching or exposed to relatively polar SDS head group environment, hence observing the decrease in the fluorescence intensity with significant red shift in emission maxima.

The change in surface hydrophobicity of PNA with addition of SDS was further explored with 8-anilino-1-naphthalenesulfonic acid (ANS), an extrinisc fluorophore to probe hydrophobic surface in protein. ANS preferentially binds to the hydrophobic patches of protein, and as per the strength of hydrophobic binding, fluorescence property of ANS changes [56]. The unbound ANS has very weak fluorescence emission in the range of 400 - 600 nm with maxima around 525 nm, upon excitation at 380 nm [56]. However upon binding to hydrophobic pocket, the emission maxima blue shifts to shorter wavelength, and the extent of shift depends upon affinity of the binding interface [57]. In intact PNA, ANS was excited at 388 nm and the emission was observed from 400 to 600 nm with the maxima centred at 525 nm and 500 a.u. intensity (Fig. 2C and 2F). The extrinsic fluorophore property indicated that the intact protein is very compact with samller number of ANS accessible hydrophobic patches (Fig. 2C). In corroboration with the tryptophan fluorescence and CD data, in presence of SDS, changes in ANS accessible hydrophobic patches is observed. The ANS fluorescence parameter increased exponentially from 750 a.u. intensity in presence of 1 mM SDS to 2500 a.u. in 20 mM SDS with maxima centered at 510 nm, i.e. a blue shift in emission maxima by 15 nm (Fig. 2F), which indicated that the surfactant brought significant changes in PNA conformation that increases the ANS accessible hydrophobic patches in the protein, i.e. SDS binding to the protein exposes the hydrophobic patches of the protein. Thus, the change in the protein-tryptophan/tyrosine fluorescnece intensity, upon addition of SDS, is due to net increase in polarity around the fluorophore. Additionally, upon correlating single ligand docking data with ANS data, it is likely that the SDS binding in second hydrophobic region (i.e. between sheet 2 and three loops) of PNA is bringing the conformational changes exposing the hydophobic region to ANS, and more significantly above the CMC of SDS.

### 3.3 CTAB induced structural perturbations in PNA

CTAB is taken as representative amphiphilic detergent with positively charged ammonium head group and 16 carbon containing alkyl chain. This cationic detergent is reported to have a CMC value around 0.2 mM in phosphate buffers, pH 7.4. CTAB is highly reactive in terms of protein unfolding, and it induces significant conformational change in protein at relatively lower concentrations than anionic detergents [58]. In the same line, the CD spectrum of PNA showed significant change in the protein ellipticity, from -3 mdeg to -11 mdeg at 205 nm, as CTAB concentration was increased from 0.2 to 2 mM (Fig. 3A and 3D). Although, the insignificant change in ellipticity was observed for 0.2 mM or lower CTAB concentrations. For CTAB concentration 0.2 to 2 mM, a shift in ellipticity minima from 220 nm to 207 nm was observed. The ellipticity trough at 207 nm was accompanied with a broader shoulder ranging from 216 nm to 223 nm (Fig.3A). Additionally, the positive ellipticity peak at 195 nm showed 4 folds decrease in the signal with no significant shift in the peak (Fig. 3A). Likewise, the ellipticity at 205 nm showed 3 folds increase, as CTAB concentration in PNA solution was increased from monomeric population to micellar population (Fig. 3D). Thus, in parallel to SDS effect above its CMC, PNA underwent conformational rearrangement, indicating presence of α-helical conformation in presence of above CMC concentration of CTAB, i.e. > 0.2 mM. Like the mapping of SDS binding interface over PNA, CTAB had also shown to dock near the SDS binding pocket, i.e. in second hydrophobic core, but with slightly lower binding energy, i.e. -2.52 KCal/mol (Table 1). The residues involved in interaction with the surfactants were Thr124, Pro134, Asp136 and Pro152 (Table1, Fig. S4-5). The charged ammonium group form polar interaction with Asp136, Thr124 side chain and Pro134, whereas Pro152 formed weaker van der Waals and non-polar interactions with alkyl side chains (Fig. S5). Interestingly, the binding interface of CTAB-PNA was in close proximity of Tyr153, which is further used as intrinsic fluorophore to observe the conformational changes in presence of the surfactant.

**Figure 3:**
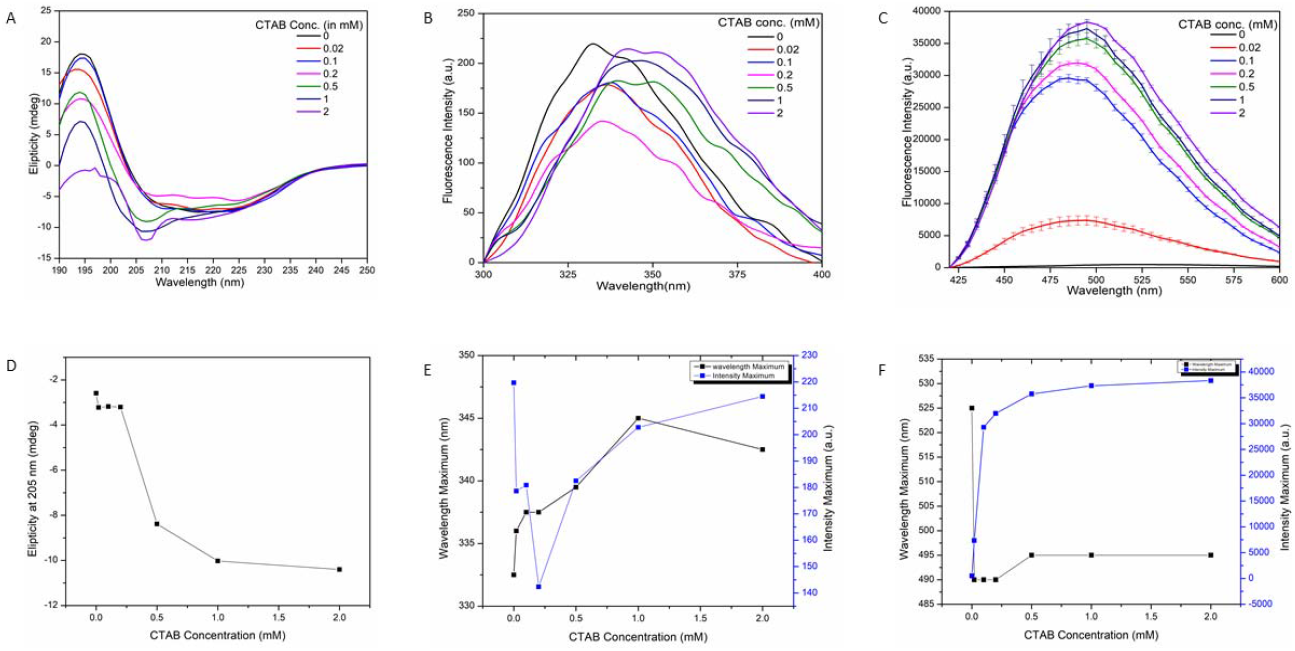
Conformational changes in PNA in presence of varying concentration of CTAB. (A) Far–UV CD spectra from 190 nm to 250nm; (B) tryptophan fluorescence emission spectra from 300 nm to 400 nm; (C) ANS fluorescence spectra from 420 nm to 600 nm upon binding of PNA at increasing concentration of CTAB; (D) ellipticity maxima at 205 nm plotted against increasing concentration of CTAB in far-UV CD spectra; (E) intensity and wavelength maximum of tryptophan fluorescence plotted against increasing concentration of CTAB; and (F) intensity and wavelength maximum of ANS fluorescence intensity plotted against increasing concentration of CTAB.

To observe the conformational rearrangement in presence of CTAB, intrinsic fluorophores of PNA were excited at 280 nm, and their emission spectra was collected in the range of 300 to 400 nm. The varying concentrations, 0 to 2 mM, of CTAB was used to study the conformational changes in PNA. PNA only solution, upon excitation at 280 nm, gave an emission peak at 332 nm with intensity of 220 a.u. (Fig. 3B). The intrinsic fluorescence intensity and emission maxima of PNA solution significantly changed upon addition of increasing CTAB concentration (Fig. 3B and 3E). The intensity first decreased to ∼140 a.u. till 0.2 mM CTAB, then it retained native like fluorescence intensity of 215 a.u. at 2 mM CTAB. However, the emission maxima showed a gradual Bathochromic shift with increasing CTAB concentration from 0 to 2 mM with plateau at ∼345 nm (Fig. 3E). Likewise, the ellipticity at 205 nm showed 3 folds increase, as CTAB concentration in PNA solution increased from monomeric population to micellar population (Fig. 3D). This indicates a possible conformational rearrangement in PNA as a result of change in local Physicochemical environment of the intrinsic fluorophore. Upon increasing the concentration of CTAB up to 0.2 mM, the indole ring of tryptophan or 4-hydroxy phenyl group of tyrosine might be getting exposed to polar environment due to conformational rearrangement in PNA or binding of polar head in atomic proximity. It has been reported that emission maxima and intensity of tryptophan/tyrosine are affected by polarity of micro-environment, hydrogen bond, exposure to incident light and other non-covalent interactions [59]. According to this literature, with increasing polarity in microenvironment of tryptophan/tyrosine, there is noticeable red shift in emission maxima. In the same line, CTAB binding to PNA gradually shifted the emission maxima to 345 nm from 330 nm (Fig. 3E), i.e. a red shift of 15 nm, whereas SDS binding showed a red shift of 24 nm (Fig. 2E). The comparatively weaker red shift in intrinsic fluorescence and ellipticity further rationalized the weaker binding energy of CTAB for the protein than of SDS.

ANS, like for SDS system, extrinsic fluorescence was used to track change in PNA hydrophobic surface upon the CTAB treatment. Lowest concentration of CTAB, i.e. 0.02 mM, brought significant changes in the protein surface, as the ANS fluorescence intensity and emission maxima shifted to 7500 a.u. at 490 nm from 500 a.u. at 525 nm, respectively (Fig. 3C and 3F). Though further addition of CTAB, till CMC, have insignificant effect on emission maxima, but the emission intensity further increased to ∼32000 a.u. for the 0.2 mM surfactant, i.e. 64 folds rise in fluorescence intensity (Fig. 3F), indicating drastic change in accessible hydrophobic surfaces of PNA. In accordance with CD and intrinsic fluorescence data, there is additional 12 folds rise in intensity with further increasing CTAB concentration from 0.2 mM to 2 mM (Fig. 3F). Above the CTAB CMC, ANS accessible hydrophobic patches increases in PNA. This increase in ANS fluorescence intensity was also accompanied by red shift in maxima from 490 nm to 495 nm. In some of the studies, SDS has been demonstrated to promote protein aggregation [50, 60-62] In previous studies, it has been found that ANS has comparatively lower affinity for native states of protein but has higher affinity for partially unfolded confirmations with accessible hydrophobic surfaces, which accordingly enhances ANS fluorescence intensity upon interaction [63]. The positively charged ammonium group of CTAB interacts with negatively charged amino acids in protein, resulting in reshuffling of native interaction network giving rise to alternative conformation with new free energy minima [64]. It has also been noticed that the alkyl chains of CTAB interacts with interior region of protein and non-polar parts via hydrophobic interactions [65]. The electrostatic and hydrophobic interactions have been found to play a crucial role in protein-surfactant interactions [64-66]. To this end, CTAB data further supports the finding that the cationic surfactant induces conformational rearrangement in PNA relatively more towards hydrophobic accessible surfaces than the anionic surfactant does.

### 3.4 Molecular Dynamics simulations for PNA-SDS/CTAB system

The root mean square deviation (RMSD) provides information about structural stability and conformational change in protein structure. Time evolution of backbone RMSD was calculated for PNA only and PNA with SDS or CTAB systems (Fig 4A). For the PNA only system, RMSD remained within 0.5 nm for the last 200 ns, however, a slight increase was observed after 450 ns. For the PNA-SDS or PNA-CTAB system, the RMSD values for the last 200 ns simulations remained relatively stable with fluctuation in the range of 0.25 - 0.4 nm (Fig 4A). To analyse the comparative overall spread of RMSD, the boxplots were done for all the three systems; in the PNA only system, the RMSD values were spread between ∼0.30 nm to ∼0.60 nm, whereas for the PNA-SDS/-CTAB systems, RMSD values were mainly spread between ∼0.24 nm to ∼0.35 nm and ∼0.10 nm to ∼ 0.47 nm, respectively (Fig. 4B). To further explore the protein residue contributing to the RMSD spread, root mean square fluctuations (RMSF) were plotted for the backbone contribution from each residue of the protein for all the three systems (Fig. 4C). The C-terminal region, 150-160, 178-182 and 95-105 residues of the protein only system was relatively highly flexible, compared to the other two systems, resulting in broader RMSD spread, especially in the last 300 ns simulation (Fig. 4C). Among the surfactants, ∼70-75, ∼90-95, ∼210-212 residues of PNA showed slightly higher fluctuations in the presence of SDS compared to CTAB system, and ∼110-115, ∼121-138, and ∼180-182 residues showed slightly higher fluctuations in CTAB system compare to SDS system. To observe overall change in the protein structure upon addition of the surfactant, radius of gyration (ROG) of PNA in the systems (PNA only and PNA-SDS/-CTAB systems) were measured (Fig. S6). The ROG showed insignificant change over the studied simulation time, i.e. 500 ns (Fig. S6A), and the boxplots of ROG showed that the radius in all the three systems fluctuated between ∼1.7 nm to ∼1.8 nm with lower values for the protein-surfactant system than the protein only system (Fig. S6B). Interestingly, the protein ROG reduced upon addition of the surfactant to the system, indicating that PNA attains further compact structure in presence of the surfactant. Overall, the data suggested that the protein, well equilibrated in all the three systems, showed relatively reduced backbone fluctuations, resulting in compact PNA structure in presence of the surfactant. The region undergoing change in fluctuation/compactness falls in proximity of the binding pockets of the surfactant as shown by spectroscopic and docking data, thus corroborating the experimental data.

**Figure 4:**
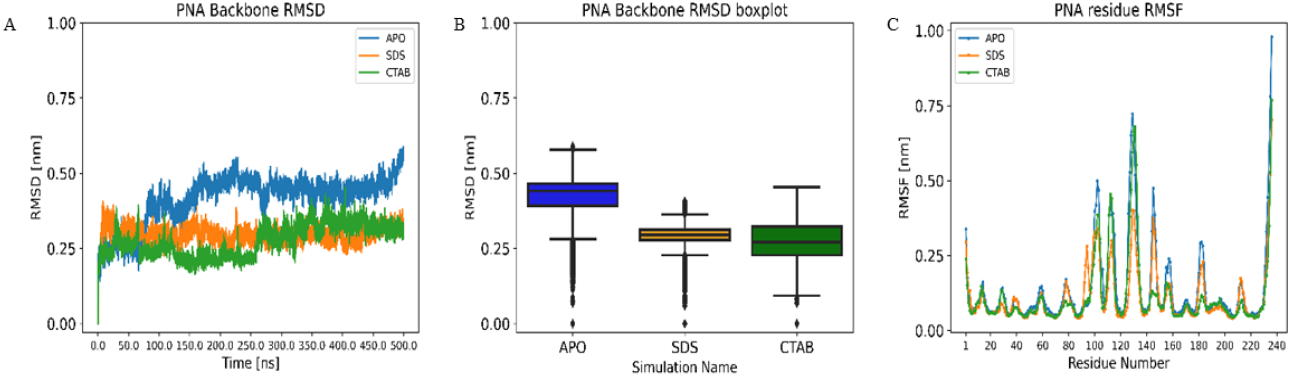
(A) Time evolution of backbone RMSD in PNA only, PNA with SDS or CTAB systems. (B) Boxplots of backbone RMSD in PNA only, PNA with SDS or CTAB systems. (C) RMSF of backbone of PNA only, PNA with SDS or CTAB systems.

Fig. 5 showed variation in the secondary structure of PNA during the 500 ns simulation run. The PNA only system consists of β-sheet and turns (Fig. 5A). Also, a minor population of α-helix and bend are found in the protein. In presence of SDS (Fig. 5B), the residues ∼1-10 mostly consist of β-sheet, which gradually changed into coil during the last 200 ns simulation. The residues ∼100-120 showed more turn and bend in SDS simulation as compared to the PNA only simulation. In residues ∼121-130, the bends are majorly converted into the coil with progression of simulation time, as seen in the case of the CTAB-PNA system (Fig. 5C), whereas not many changes were seen for the residues in SDS system (Fig. 5B). Additionally, the residues from ∼200-220 in case of both the surfactant system, SDS and CTAB systems, showed more bend from 50 – 500 ns in comparison to turns in PNA only system. However, in the case of CTAB simulation, ∼181-200 residues showed conformational transition from α-helix to turn within the 50-450 ns simulation time frame.

**Figure 5:**
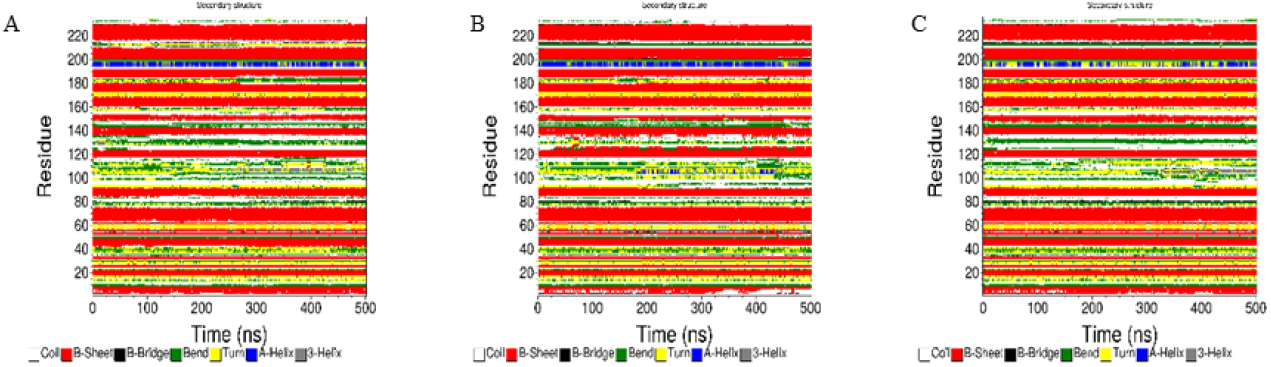
(A) Time evolution of secondary structure changes in PNA in Apo form. (B) in the presence of SDS molecules. (C) in the presence of CTAB molecules.

Fig. 6 (A and B) showed the average minimum distance between each residue of PNA from each SDS molecule. The average minimum distance between PNA residues regions ∼1-21, ∼47-51 ∼161-200, and ∼221-230 remained close (< 0.6 nm) to SDS throughout the simulation time (Fig. 6B). Major interacting amino acids were Pro16, Met 55, Leu 116, Thr 142, Ile 197 and Ile 199 residues for at least 50% or more simulation time (Fig. 6B-C). This data further corroborates molecular docking (Table 1) and intrinsic fluorescence data (Fig. 2B), which indicated tryptophan/tyrosine fluorescence quenching with increasing concentration of SDS is either for the local physicochemical conditions, atleast around Trp55 residue, changes towards net higher polarity or progressive presence of hydrophobic functional groups such as Pro16, Leu66, Val142, Ile166, Pro199 and SDS hydrophobic tail are quenching the fluorescence, or both. In PNA-CTAB system, residues from ∼1-11 and ∼101-145 remained close to CTAB molecules throughout the simulation time (Fig. 6D-E). PNA residues Leu 66, Thr 105, Tyr 124, His 136, Val 153, Thr 164, and Ile 166 interacted with CTAB residues for at least 50% or more simulations time. This data corroborates with intrinsic fluorescence data (Fig. 3B) showed that upon increasing the concentration of CTAB up to 0.2 mM, 4-hydroxy phenyl groups of Tyr 130 and Tyr 153 are being exposed to polar environment due to conformational rearrangement in PNA or binding of polar head group in atomic proximity.

**Figure 6:**
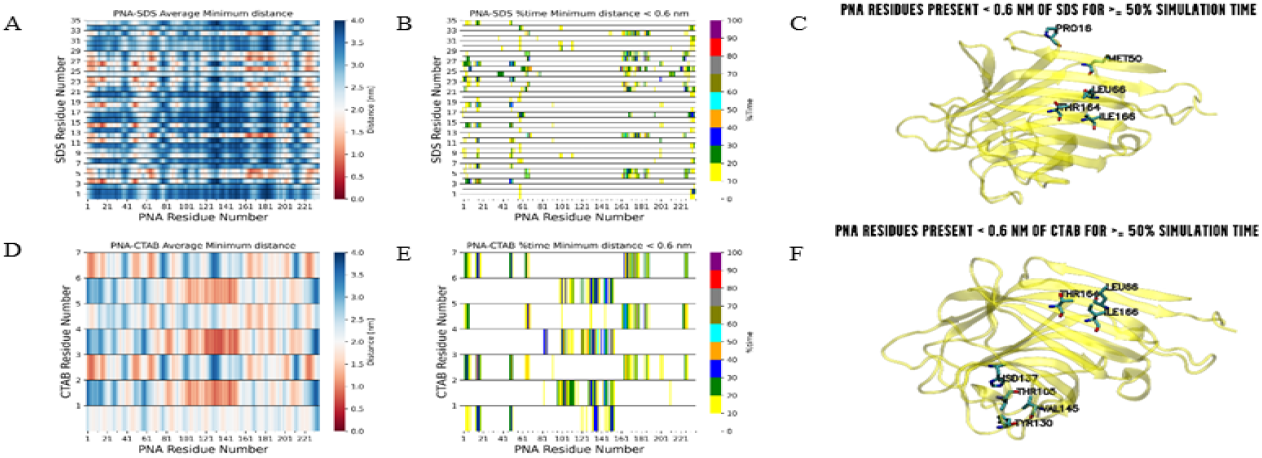
(A) Average minimum distances between PNA residues and SDS residues. (B) Percentage of time PNA residues remained within < 0.6 nm of SDS molecules. (C) Residues that remained >= 50% simulations time < 0.6 nm of SDS molecules. (D) Average minimum distances between PNA residues and CTAB residues. (E) Percentage of time PNA residues remained within < 0.6 nm of CTAB molecules. (F) Residues that remained > = 50% simulations time < 0.6 nm of CTAB molecules.

To further investigate the interfacial interactions in PNA-SDS or PNA-CTAB system, the time evolution of interaction energy and its components (van der Waals and electrostatic energy) were calculated. The data showed that the multiple SDS molecules (∼25 no.) interacted with PNA predominantly through van der Waals interaction than electrostatic (Fig. 7A-B). Whereas, in defined box, 3-4 CTAB molecules showed to interact with PNA predominantly through electrostatic interaction than van der Waals interaction (Fig. 7D-E). Taking both the interactions together for both the system, the time evolution interaction energy showed that multiple CTAB molecules interacted more strongly with PNA, than multiple SDS molecules (more negative value of total interaction energy, Fig. 7C and 7F).

**Figure 7:**
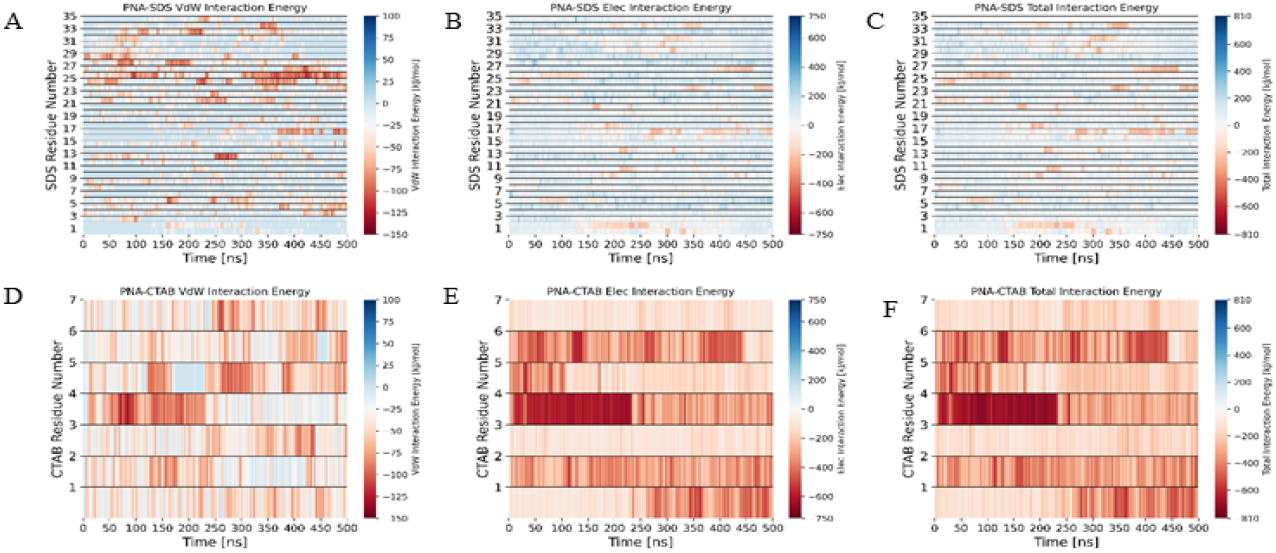
(A) Time evolution of VdW interaction energy between PNA and SDS molecules. (B) Time evolution of Electrostatic interaction energy between PNA and SDS molecules. (C) Time evolution of total interaction energy between PNA and SDS molecules. (D) Time evolution of VdW interaction energy between PNA and CTAB molecules. (E) Time evolution of Electrostatic interaction energy between PNA and CTAB molecules. (F) Time evolution of total interaction energy between PNA and CTAB molecules.

## CONCLUSION

The interactions of PNA with a cationic detergent, CTAB, and an anionic detergent, SDS, were explored. The data suggested that SDS induces α-helices in PNA, above its CMC, which in consistence with the findings reported by Khan et al. for SDS effect on another lectin binding protein ConA [50]. Other studies have also depicted protein denaturation by SDS using different proteins such as ubiquitin and lysozyme [67, 68]. MD simulations data shows that the major interaction between the SDS micelle and PNA is VdW interactions, which are weak and dynamic in nature, and could be caused due to the spatio-temporal arrangement on the ligands in the SDS-PNA complex. In general, it is proposed that SDS denatures proteins by forming protein-decorated micelles (core-shell model). Depending on the SDS:protein ratio and the protein molecular mass, the formed structures can range from multiple partly unfolded protein molecules surrounding a single shared micelle to a single polypeptide chain decorating multiple micelles [10]. On the other hand, CTAB has a CMC below SDS, i.e. 1 mM [69], and was shown to disrupt the PNA structure, at much lower concentration. On the contrary to SDS interaction mode, the predominant interactions between CTAB and PNA are electrostatic interactions. CTAB has shown to disrupt ConA protein structure, and led lysozyme protein to form amyloid structure [11, 70]. Additionally, the spectroscopic data has shown that the unfolding and refolding of human serum albumin occur with increasing CTAB concentration from its monomeric form to micellar form, i.e. from below to above its CMC [71]. Taking spectroscopic, docking and simulation data altogether, it is concluded that the surfactant of oppositely charged head group, irrespective of the binding stoichiometry, induces conformational rearrangement in PNA around its monomeric to micellar phase boundary, where the protein attains compact structure with more hydrophobic accessible surfaces.

## Credit Authorship Contribution Statement

**Dr. Shreyasi Asthana** has conceptualised the work, and executed most of the *in vitro* experiments alon with **Ms. Sonali Mohanty**, who has also helped SA with writing the rough draft and the figures compiling. **Mr. Harshit Kalra** and **Ms. Nandini Karunanethi** have helped with the ANS fluorescence and docking data respectively. **Dr. Sujit Kumar** has helped us with the data analysis. **Dr. Nikhil Agrawal** has run all the molecule dynamics simulations, and helped **Dr. Suman Jha** finalising the manuscript in current format. **Dr. Suman Jha** and **Dr. Nikhil Agrawal**, have, additionally, acquired the fund for the work, conceptualised the work, coordinated the in vitro and *in silico* measurements, administered and supervised the project.

## Acknowledgments

S.J. acknowledges the financial support from the Department of Science and Technology (ST-BT-MISC-0020-2018-4002/ST), Govt. of Odisha, India and Indian Council of Medical Research (35/05/2020-Nano/BMS), govt. of India, India for the study. Additionally, N.A. acknowledges the ERDF grant (No. 1.1.1.2/VIAA/4/20/757) for financial support, and thanks the Centre for High Performance Computing (CHPC) in Cape Town (South Africa) for supercomputing resources.

## Supporting Information

The supporting information files contain PDB structure of PNA, docking results and ROG of the three system, and has been attached.

